# Monitoring RNA restructuring in a human cell-free extract reveals eIF4A-dependent and eIF4A-independent unwinding activity

**DOI:** 10.1101/2022.12.14.520460

**Authors:** Mattie H. O’Sullivan, Christopher S. Fraser

## Abstract

The canonical DEAD-box helicase, eIF4A, unwinds 5’ UTR secondary structures to promote mRNA translation initiation. Growing evidence has indicated that other helicases, such as DHX29 and DDX3/ded1p, also function to promote the scanning of the 40S subunit on highly structured mRNAs. It is unknown how the relative contributions of eIF4A and other helicases regulate duplex unwinding on an mRNA to promote initiation. Here, we have adapted a real-time fluorescent duplex unwinding assay to precisely monitor helicase activity in the 5’ UTR of a reporter mRNA that can be translated in a cell-free extract in parallel. We monitored the rate of 5’ UTR-dependent duplex unwinding in the absence or presence of Hippuristanol, a dominant negative eIF4A (eIF4A-R362Q), or a mutant eIF4E (eIF4E-W73L) that can bind the m^7^G cap but not eIF4G. Our experiments reveal that roughly 50% of the duplex unwinding activity in the cell-free extract can be attributed to an eIF4A-dependent mechanism, while the remaining 50% of duplex unwinding activity is attributed to an eIF4A-independent mechanism. Importantly, we show that the robust eIF4A-independent duplex unwinding is not sufficient for translation. We also show that the m^7^G cap structure, and not the poly(A) tail, is the primary mRNA modification responsible for promoting duplex unwinding in our cell-free extract system. Overall, the fluorescent duplex unwinding assay provides a precise method to investigate how eIF4A-dependent and eIF4A-independent helicase activity regulates translation initiation in cell-free extracts. We anticipate that potential small molecule inhibitors could be tested for helicase inhibition using this duplex unwinding assay.

## INTRODUCTION

Eukaryotic initiation factors unwind mRNA secondary structure so that a single-stranded region can be positioned directly in the mRNA binding channel of the 40S ribosomal subunit. Eukaryotic initiation factor (eIF) 4A is the DEAD-box helicase that has been proposed to directly unwind mRNA secondary structure. As a member of the helicase superfamily 2 (SF2), eIF4A possesses two RecA-like domains that enable adenosine triphosphate (ATP) binding and hydrolysis, as well as RNA binding and duplex unwinding (1,2). To regulate its helicase activity, eIF4A can be associated with several accessory proteins, including eIF4G, eIF4B, and eIF4H (reviewed in (3,4)). These accessory proteins function to activate the duplex unwinding activity of eIF4A, enhance its RNA binding activity, and accelerate the opening and closing of its RecA-like domains (5–11). The binding of eIF4G and eIF4B/H transform eIF4A from a non-processive helicase to a processive helicase (12). For translation initiation, eIF4A is positioned at the mRNA exit channel of the 40S ribosomal subunit, thereby promoting slotting of the 5’ untranslated region (5’ UTR) into the mRNA binding channel and decoding site of the 40S subunit (13). The eIF4A helicase can exist as a component of the eIF4F complex or in free form (14). Importantly, the helicase activity of eIF4A is required for the translation of all mRNAs regardless of their structural complexity (15).

In addition to eIF4A, other helicase proteins have been associated with removing secondary structure in the 5’ UTR of mRNA. These include DHX29 and DDX3/ded1p (reviewed in (3)). These helicases have been reported to bind to eIF4G and/or the 40S ribosomal subunit, providing a possible mechanism for their specific recruitment to the initiation complex (16–21). Both DDX3/ded1p and DHX29 have been implicated in regulating specific mRNA translation, especially mRNAs that have 5’ UTRs with the potential to contain extensive secondary structure (22–24). It is possible that additional helicase proteins beyond eIF4A, DDX3/dedp1, and DHX29 may regulate the degree to which secondary structure may form in mRNAs. Monitoring global RNA structure has revealed that there are substantially fewer structured mRNA regions found in live yeast and mammalian cells than expected (25). This finding was largely attributed to energy-dependent processes, such as RNA helicases. Interestingly, it was proposed that this helicase activity is not dependent on the process of translation since secondary structure is absent from both coding and non-coding regions of mRNA (25). Thus, it is possible that RNA helicases play a role independently of the ribosome to regulate the amount of secondary structure found in mRNAs. Consistent with this model, eIF4A has been proposed to function as a non-specific ATP-dependent RNA chaperone that limits the condensation of RNA (26). This chaperone-like activity is independent of its activity in eIF4F and translation initiation, perhaps explaining why the cellular concentration of this protein is in a large excess compared to the translation machinery.

The duplex unwinding activity of various helicase proteins have been successfully characterized using highly purified components in ensemble and single molecule assays. While very informative, using purified reconstituted systems to study helicase proteins is generally limited by the fact that they may not include all helicase proteins needed for a process, or all the accessory factors that are important for their regulation. Thus, it would be valuable to precisely monitor duplex unwinding in a more complete system to better understanding of how multiple helicases and their accessory proteins may function together to remove secondary structure in mRNAs. To this end, we reasoned that such a system could be provided by a cell-free cytoplasmic extract, which has been a powerful system to study various aspects of translational control over many decades. We previously established an *in vitro* fluorescence assay able to detect RNA strand separation using purified recombinant eukaryotic initiation factors (eIFs) (5,27,28). The assay uses two short modified RNA reporter strands designed to anneal to engineered binding sites on an unmodified RNA loading strand. A robust increase in cyanine 3 (Cy3) fluorescence is observed when one strand is separated from the loading strand and its closely annealed quencher dye-modified strand. Using this approach, we have analyzed duplex unwinding on ~1 Kb loading strands that contain internal ribosome entry sites (IRESes)(29,30). A similar fluorescent duplex unwinding assay has been used to monitor the helicase activity on a very short RNA substrate using cell-free extracts of marine organisms (31). While that fluorescent assay is incompatible with monitoring duplex unwinding of a full-length mRNA it did suggest that helicase activity can be monitored in a cell-free extract system.

Here, we have optimized a real-time fluorescent duplex unwinding assay to precisely monitor duplex unwinding in the 5’ UTR of a full-length mRNA reporter in a HeLa cell-free extract. In addition to displaying high fidelity translation that is m^7^G cap and poly(A) dependent, our HeLa cell-free extract exhibits a robust ATPase dependent duplex unwinding activity. Using Hippuristanol or a dominant negative mutant of eIF4A to specifically inhibit eIF4A-dependent helicase activity, we demonstrate that HeLa cell-free extracts possess roughly 50% eIF4A-dependent and 50% eIF4A-independent duplex unwinding activity. We further show that a mutant eIF4E, eIF4E-W73L, also reveals a substantial amount of eIF4F-independent duplex unwinding in HeLa cell-free extracts. The m^7^G cap and poly(A) tail function synergistically to regulate mRNA translation in live cells and cell-free extracts (32–35). We therefore used our fluorescent duplex unwinding assay to demonstrate that the m^7^G cap structure, and not the poly(A) tail, is the primary mRNA modification responsible for promoting duplex unwinding in the cell-free extract system. Overall, our fluorescent duplex unwinding assay reveals how eIF4A-dependent and eIF4A-independent helicase activity is present in cell-free extracts and provides a robust method to further investigate these activities.

## RESULTS

### Monitoring RNA duplex unwinding in nuclease-treated cell-free extracts

To monitor duplex unwinding of a translationally active mRNA, we engineered fluorescently-modified reporter strand annealing sites into the 5’ UTR of a mRNA that codes for the bioluminescent gene, Nano Luciferase (NLuc) (Fig. 1). The *in vitro* T7 transcribed mRNA possesses both a m^7^G cap and poly(A) tail so that it can be faithfully translated when added to a cell-free cytoplasmic extract from HeLa cells (Fig. 1). A 50 nt poly(A) tail was chosen due to it being reported as an appropriate length required for efficient PABP binding (36). Cell-free extracts generated from HeLa S3 suspension cells display a linear rate of translation for a 50 nM concentration of the NLuc reporter mRNA, which includes annealed reporter strands, for 30 mins at 30 °C (Fig. 2A and S1). Linear translation for 30 mins indicates that the annealed reporter strands do not promote mRNA degradation during the time course of the experiment. We note that the rate of translation is not appreciably inhibited by the addition of the annealed beacons (Fig. S2). Cell-free extracts from mammalian cells typically display faithful synergism between the m^7^G cap and poly(A) tail for translation (37–39). To verify this is true for our cell-free extracts, we generated combinations of mRNAs that possess either a m^7^G cap, poly(A) tail, or both. Monitoring luciferase after 30 mins incubation at 30 °C reveals a strong dependence of the m^7^G cap structure for translation and a roughly twofold synergy between the m^7^G cap and poly(A) tail (Fig. 2B).

**Figure 1.**
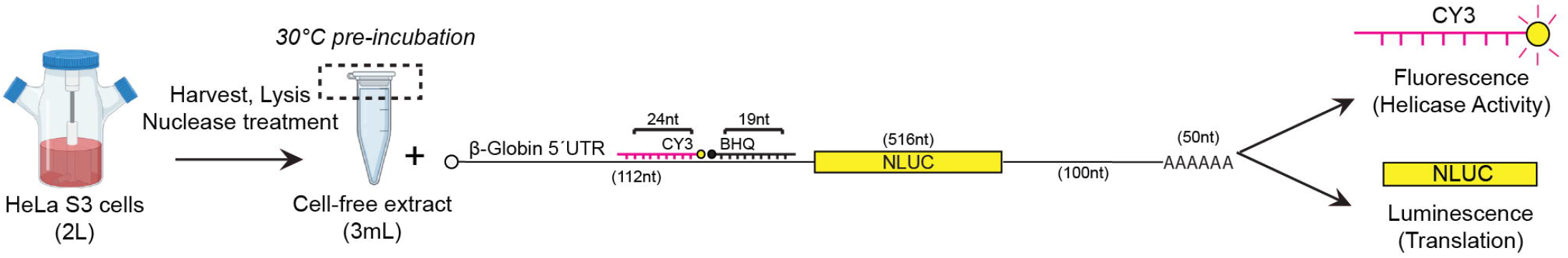
Schematic of the development of cell-free extracts for use with a mRNA reporter to monitor unwinding of RNA duplexes in parallel with changes in protein synthesis. Cell-free extracts are generated from HeLa S3 cells grown in large liquid suspension cultures (2 liters). A modular full-length mRNA reporter is used to monitor: a) duplex unwinding via fluorescent beacons annealed to the mRNA 5’ UTR; and b) translation of a bioluminescent gene (NLuc). Lysates are preincubated for 10 min at 30 °C to allow any supplemented proteins or inhibitors to reach equilibrium prior to adding the mRNA reporter and recording assay measurements. Suspension cell flask depicted in this figure was created with BioRender.com.

**Figure 2.**
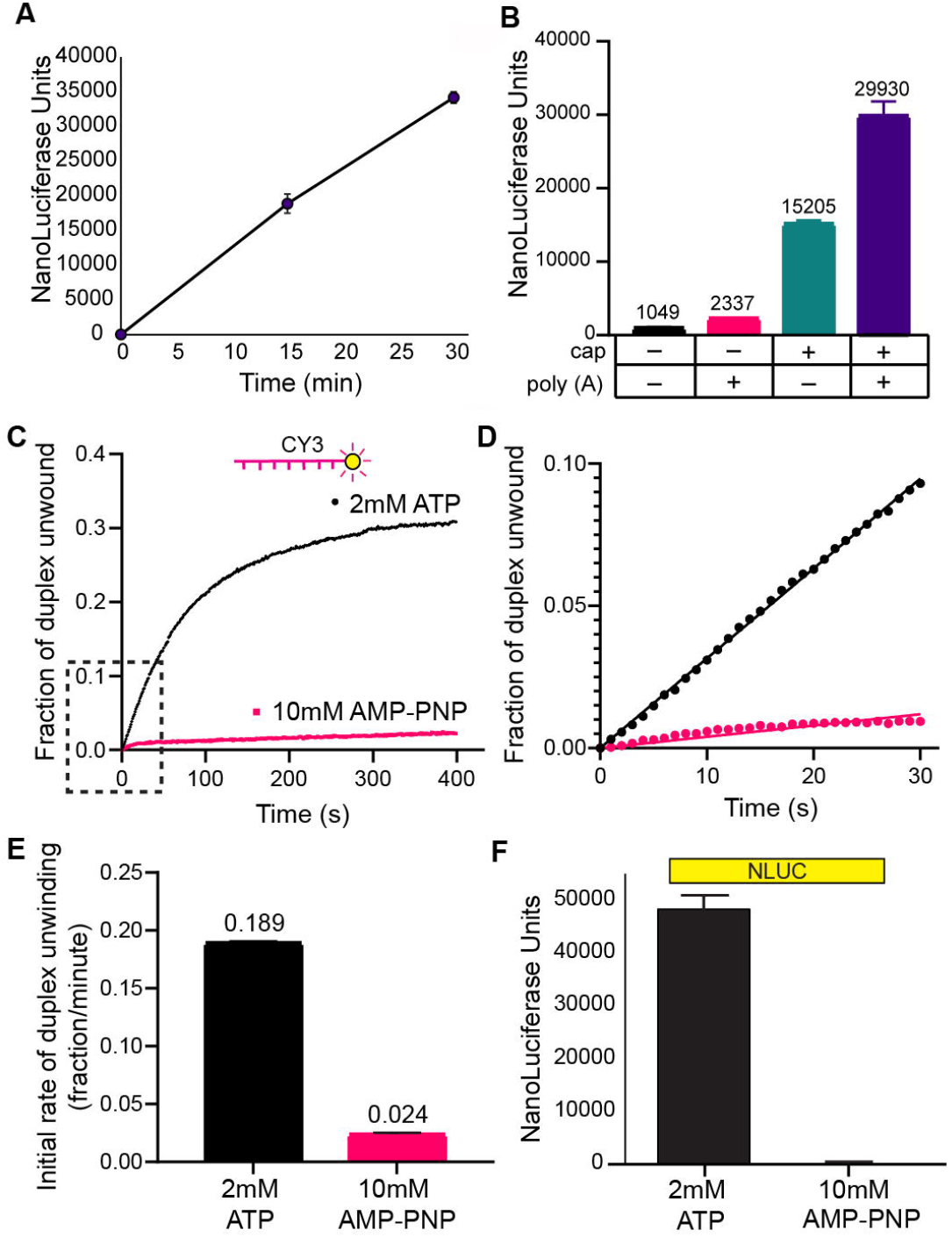
RNA duplex unwinding in nuclease-treated cell-free extracts is ATP dependent. (**A**) Time course of translation in cell-free extracts using the reporter depicted in Fig. 1. Lysate and translation assay mix was pre-incubated for 10 min at 30 °C prior to adding the mRNA (50 nM) to initiate protein synthesis for the time shown, as described in *Experimental Procedures*. (**B**) Bar graph depicting luciferase translation of mRNA reporter(s) with each combination of m^7^GTP and poly(A) tail modifications to the mRNA, measured after 30 min incubated at 30 °C. (**C**) The average trace of three independent time courses is shown for the fraction of duplex unwound in cell-free extracts in the presence of 2 mM ATP (black) or 10 mM AMP-PNP (magenta). The zero time point represents the fraction of duplex unwound immediately upon mRNA addition to the cell-free extract, as described in *Experimental Procedures*. A change in total Cy3 fluorescence is converted into the fraction of duplex unwound versus time after ATP/AMP-PNP addition, as described in the *Experimental Procedures*. (**D**) A magnified view of the dashed region of the unwinding time course (C) is shown. Initial rates of the unwinding time course are determined by linear fits to the linear portion of the data. (**E**) The initial rate of duplex unwinding (fraction unwound per min) by each dataset are shown as indicated. (**F**) NLuc translation was measured in cell-free extracts over 30 min. Error bars for all data indicate the standard errors of the mean.

To study duplex unwinding associated with the mRNA reporter, we monitored the change in Cy3 fluorescence in real time. Accordingly, we preincubated the HeLa cell extract with 2 mM ATP and all components minus the mRNA reporter for 10 mins at 30 °C. Upon addition of the mRNA reporter (50 nM), we observe a robust increase in Cy3 fluorescence that is converted to the fraction of duplex unwound (Fig. 2C), as described in the *Experimental Procedures*. The initial rate of duplex unwinding is calculated from a linear fit to the initial linear portion of the time course, as described in the *Experimental Procedure*s (Fig. 2D). We observe an initial fraction of duplex unwound per min of 0.189 ± 0.001 (Fig. 2E and Table 1). To verify that the change in Cy3 fluorescence is due to authentic ATPase activity in the cell-free extract, we preincubated the HeLa cell extract with 10 mM of the non-hydrolysable ATP analog, AMP-PNP, for 10 mins at 30 °C. Upon addition of the mRNA reporter, a dramatic reduction in the change in Cy3 fluorescence over time is observed in the presence of 10 mM AMP-PNP (Fig. 2C). The initial rate of duplex unwinding shows that the initial fraction of duplex unwound per minute in the presence of 10 mM AMP-PNP is 0.024 ± 0.001 (Fig. 2D, 2E and Table 1). This allows us to calibrate duplex unwinding activity in our cell-free extracts and shows that a roughly 8-fold change in unwinding activity is observed between active (ATP) and inactive (AMP-PNP) conditions. Consistent with the inhibition of Cy3 fluorescence in the presence of 10 mM AMP-PNP, we observe a dramatic inhibition of luciferase translation after 30 mins in a parallel reaction with identical reaction conditions (Fig. 2F). Together, these data reveal that translation (luciferase) and duplex unwinding (Cy3 fluorescence) can be precisely monitored in parallel in HeLa cell-free extracts.

### RNA duplex unwinding in nuclease-treated cell-free extracts is both eIF4A-dependent and eIF4A-independent

To determine what proportion of duplex unwinding in the HeLa cell-free extract is attributable to the ATPase activity of eIF4A, we tested eIF4A-specific inhibitors using our fluorescent helicase assay. Hippuristanol is a small molecule inhibitor of eIF4A that binds to and inhibits the RNA binding activity of eIF4A (Fig. 3A) (40). This drug inhibits ~70% of the helicase activity of eIF4A at a 3–5 μM concentration using our fluorescent helicase assay with purified initiation factors (28,29). Hippuristanol inhibits cap-dependent translation in cell-free extracts by ~60% even at the modest concentration of 0.4 μM (40). To determine the effect of adding Hippuristanol on helicase activity in our cell-free extract, we added 3 μM Hippuristanol (in 0.3% DMSO). In the absence of Hippuristanol, we observe a robust increase in the fraction of duplex unwound over time upon addition of the mRNA reporter in the presence of vehicle (0.3% DMSO; Fig. 3B; black line). The initial rate of duplex unwinding per min in the presence of vehicle is calculated to be 0.162 ± 0.002 (Fig. 3C, 3D and Table 1). This is similar to the rate of duplex unwinding in the absence of vehicle (Fig. 2 and Table 1). In the presence of 3 μM Hippuristanol, we observe an appreciable reduction in the change in Cy3 fluorescence over time (Fig. 3B). Calculating the initial rate of duplex unwinding in the presence of drug shows that the initial fraction of duplex unwound per minute is reduced to 0.104 ± 0.001 (Fig. 3C, 3D and Table 1). Using the initial fraction of duplex unwound in the presence of AMP-PNP to calibrate duplex unwinding, our data show that inhibiting eIF4A with 3 μM Hippuristanol reduces the rate of duplex unwinding in the 5’ UTR of the mRNA reporter by roughly 40%. Consistent with the inhibition of Cy3 fluorescence in the presence of 3 μM Hippuristanol, we observe complete inhibition of luciferase translation after 30 mins in a parallel reaction with identical reaction conditions (Fig. 3E and Table 1). We note that Hippuristanol is stored as a 1 mM stock in DMSO (*Experimental Procedures*). One is therefore limited in the concentration that can be added to a cell-free extract to inhibit eIF4A helicase activity because translation in cell-free extracts is sensitive to DMSO. Although 3 μM Hippuristanol may not completely inhibit eIF4A helicase activity, we are unable to further increase the concentration of Hippuristanol in our experiment due to the finding that a higher concentration of DMSO alone inhibits the translation activity of our reporter (Fig. 3E).

**Figure 3.**
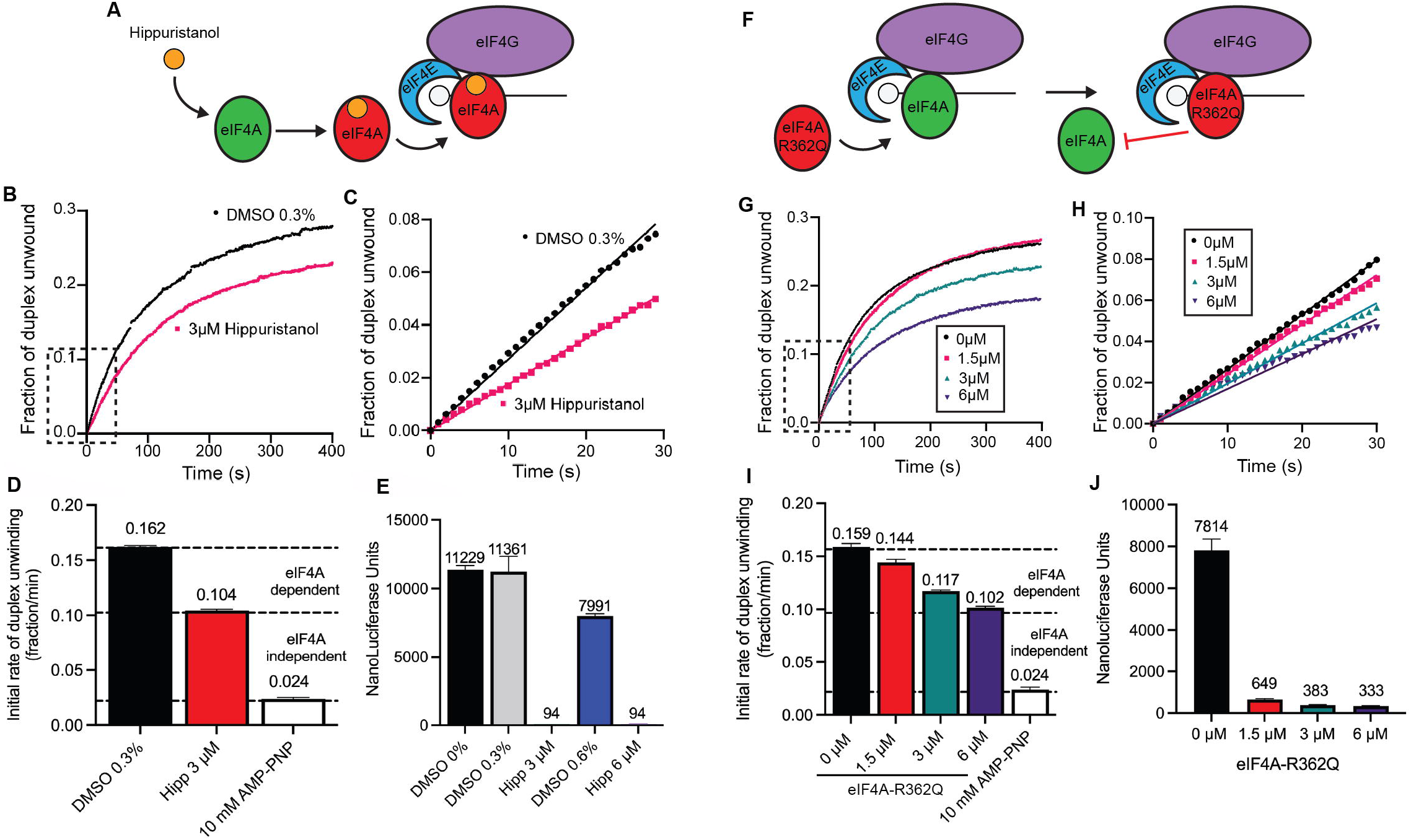
RNA duplex unwinding in nuclease-treated cell-free extracts is both eIF4A-dependent and eIF4A-independent. **(A)** Cartoon depiction of the inhibition of eIF4A by the small molecule inhibitor, Hippuristanol. While not shown for clarity, mRNAs used possess m^7^G cap and poly(A) tail **(B)** The average trace of three independent time courses is shown for the fraction of duplex unwound in cell-free extracts in the presence of vehicle (0.3% DMSO; black line) or 3 μM Hippuristanol (in 0.3% DMSO; red line). **(C)** A magnified view of the dashed region of the unwinding time course (b) is shown. Linear fits are made to the initial portion of the unwinding data in the presence of 0.3% DMSO (final concentration) ± 3 μM Hippuristanol. **(D)** The initial rates of duplex unwinding (fraction unwound per min) determined by the slope of the line in (C). A bar graph of the initial rate of duplex unwinding (fraction unwound per min) in the absence or presence of Hippuristanol or AMP-PNP is shown as indicated. Dashed lines provide an estimate of eIF4A-dependent and eIF4A-independent duplex unwinding, as determined by the rate of duplex unwinding in the absence of Hippuristanol versus the presence of Hippuristanol and AMP-PNP. The AMP-PNP value to show the calibration for ATPase-dependent duplex unwinding is reproduced from Fig. 2. **(E)** Bar graph depicting luciferase translation of mRNA reporter containing increasing concentrations of DMSO (0, 0.3 and 0.6%), and 3 μM or 6 μM Hippuristanol (diluted in 0.3% and 0.6% DMSO respectively). Luciferase activity was measured after the reporter had been incubated with the lysate assay mix for 30 min at 30 °C. (**F**) Cartoon depiction of the inhibition of eIF4A by the dominant negative eIF4A, eIF4A-R362Q. (**G**) The average trace of three independent time courses is shown for the fraction of duplex unwound in cell-free extracts in the absence (black line) or presence of eIF4A-R362Q: 1.5 μM, 3 μM or 6 μM as shown. **(H)** A magnified view of the dashed region of the unwinding time course (G) is shown. Linear fits to the initial portion of the unwinding data in the presence of 0 μM, 1.5 μM, 3 μM or 6 μM eIF4A-R362Q. **(I)** A bar graph of the initial rate of duplex unwinding (fraction unwound per min) in the absence or presence of eIF4A-R362Q and AMP-PNP is shown as indicated. Dashed lines provide an estimate of eIF4A-dependent and eIF4A-independent duplex unwinding, as determined by the rate of duplex unwinding in the absence of eIF4A-R362Q versus the presence of eIF4A-R362Q and AMP-PNP. The AMP-PNP value to show the calibration for ATPase-dependent duplex unwinding is reproduced from Fig. 2. **(J)** Bar graph depicting luciferase translation of mRNA reporter containing increasing concentrations of eIF4A-R362Q. Error bars for all data indicate the standard errors of the mean.

To further investigate the role of eIF4A in duplex unwinding in our assay, we added a recombinantly purified eIF4A-R362Q protein to the cell-free extract system. This eIF4A mutant is a well characterized dominant negative mutant that inhibits translation by preventing the recycling of eIF4A through the eIF4F complex (Fig. 3F) (41). Compared to the absence of eIF4A-R362Q, we observe a dose-dependent reduction in the change in Cy3 fluorescence over time as an increasing concentration of eIF4A-R362Q is added to the cell-free extract (Fig. 3G). Calculating the initial rate of duplex unwinding shows that the initial fraction of duplex unwound per minute is reduced from 0.159 ± 0.003 in the absence of eIF4A-R362Q to 0.102 ± 0.001 in the presence of 6 μM eIF4A-R362Q (Fig. 3H, 3I and Table 1). Using the initial fraction of duplex unwound in the presence of AMP-PNP to calibrate duplex unwinding, our data show that inhibiting eIF4A by 6 μM eIF4A-R362Q reduces the rate of duplex unwinding in the 5’ UTR of the mRNA reporter by roughly 40%. Consistent with the inhibition of Cy3 fluorescence in the presence of increasing concentrations of eIF4A-R362Q, we observe a strong reduction in luciferase translation after 30 mins in a parallel reaction with identical reaction conditions (Fig. 3J and Table 1).

These data reveal that inhibiting eIF4A activity by the addition of Hippuristanol or eIF4A-R362Q results in a modest ~40% inhibition of helicase activity in the cell-free extract. We interpret this unwinding activity to represent the eIF4A-dependent helicase activity in the cell-free extract. Importantly, a substantial amount of helicase activity remains in the presence of these eIF4A-specific inhibitors, which we interpret to represent the eIF4A-independent helicase activity in the cell-free extract. The eIF4A-independent helicase activity is not sufficient for translation, since the rate of protein synthesis is dramatically inhibited despite 60% of the helicase activity remaining in the cell-free extract. Moreover, we observe that in the presence of 1.5 μM eIF4A-R362Q, translation is strongly inhibited while the duplex unwinding activity monitored in the 5’ UTR of the reporter mRNA is inhibited by only ~10% (Fig. 3I and Table 1). These data are consistent with the finding that eIF4A is required for the translation of all mRNAs regardless of their structural complexity (15).

### RNA duplex unwinding in nuclease-treated cell-free extracts is eIF4F-dependent and eIF4F-independent

The DDX3/ded1p and DHX29 helicases have been reported to bind to eIF4G and/or the 40S ribosomal subunit (16–21). While the precise mechanism by which these helicases regulate translation is unclear, their ability to unwind secondary structure in the 5’ UTR of mRNA is likely an important part of their activity. The eIF4A-specific inhibitors used above to prevent duplex unwinding may not prevent eIF4F binding to mRNA. Moreover, Hippuristanol and eIF4A-R362Q may not prevent the recruitment of the 40S subunit since ATP hydrolysis by eIF4A is not required for mRNA recruitment to the 40S subunit mRNA binding channel (42). We therefore wanted to determine if complete inhibition of mRNA recruitment to eIF4F and the 40S subunit would result in similar inhibition of cell-free extract helicase activity compared to eIF4A inhibition alone. To this end, we generated an eIF4E protein, eIF4E-W73L, that is expected to function as an inhibitor by blocking access of the translational machinery to the 5’ m^7^G cap (Fig. 4A). Specifically, this eIF4E mutation is based on the fact that W73 in human eIF4E is required for stable binding of eIF4E to eIF4G (43,44).

**Figure 4.**
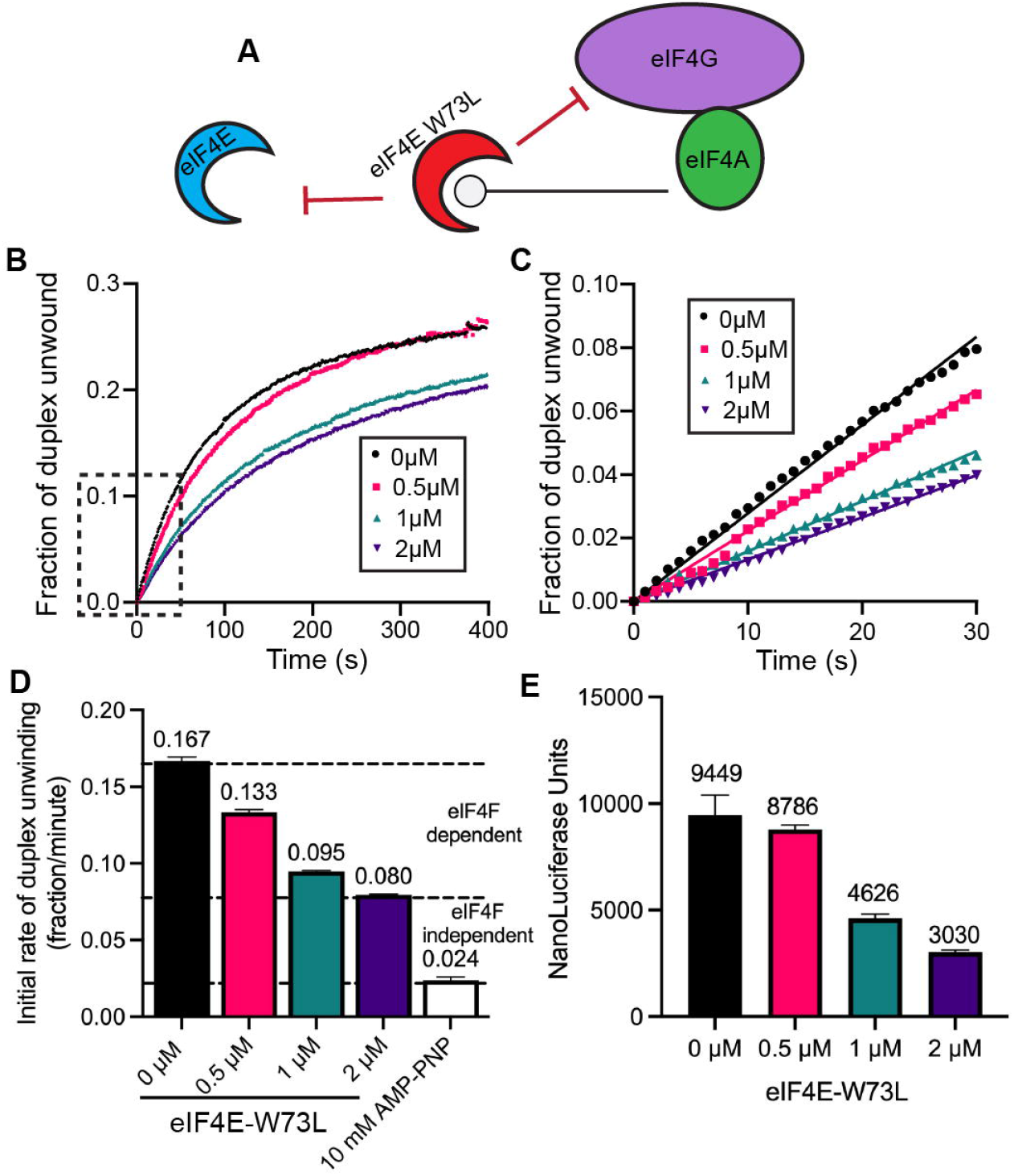
RNA duplex unwinding in nuclease-treated cell-free extracts is eIF4F-dependent. **(A)** Cartoon depicting the mechanism of translation inhibition by the dominant negative eIF4E, eIF4E-W73L. While not shown for clarity, mRNAs used possess m^7^G cap and poly(A) tail **(B)** The average trace of three independent time courses is shown for the fraction of duplex unwound in cell-free extracts in the absence or presence of increasing concentrations of eIF4E-W73L: 0 μM, 0.5 μM, 1 μM or 2 μM as shown. **(C)** A magnified view of the dashed region of the unwinding time course (B) is shown. Linear fits are made to the initial portion of the unwinding data in the absence or presence of increasing concentrations of eIF4E-W73L. **(D)** The initial rates of duplex unwinding (fraction unwound per min) determined by slope of the line in (C). Dashed lines provide an estimate of eIF4F-dependent and eIF4F-independent duplex unwinding, as determined by the rate of duplex unwinding in the absence of eIF4E-W73L versus the presence of Hippuristanol and AMP-PNP. The AMP-PNP value to show the calibration for ATPase-dependent duplex unwinding is reproduced from Fig. 2. **(E)** Bar graph depicting luciferase translation of a mRNA reporter containing increasing concentrations of eIF4E-W73L, as indicated. Error bars for all data indicate the standard errors of the mean.

To determine the effect of adding eIF4E-W73L on helicase activity, we preincubated the cell-free extract with an increasing concentration of purified protein. Compared to the absence of eIF4E-W73L, a dose-dependent reduction in the change in Cy3 fluorescence and initial rate of unwinding is observed over time as eIF4E-W73L is added to the cell-free extract (Fig. 4B and 4C). Calculating the initial rate of duplex unwinding shows that the initial fraction of duplex unwound per minute is reduced from 0.167 ± 0.003 in the absence of eIF4E-W73L to 0.080 ± 0.001 in the presence of 2 μM eIF4E-W73L (Fig. 4D and Table 1). Using the initial fraction of duplex unwound in the presence of AMP-PNP to calibrate duplex unwinding, our data show that inhibiting eIF4F recruitment by adding 2 μM eIF4E-W73L reduces the rate of duplex unwinding in the 5’ UTR of the mRNA reporter by roughly 45%. This reduction is similar in magnitude to Hippuristanol and eIF4A-R362Q. Nevertheless, we do notice that inhibiting helicase activity in the cell-free extract with eIF4E-W73L does appear to be slightly more potent than Hippuristanol and eIF4A-R362Q (Table 1). Consistent with the inhibition of Cy3 fluorescence in the presence of increasing concentrations of eIF4E-W73L, we observe a strong dose dependent reduction in luciferase translation after 30 mins in a parallel reaction with identical reaction conditions (Fig. 4E and Table 1). We note that the inhibition of duplex unwinding and translation by eIF4E-W73L appear to closely match each other in a dose dependent manner (Fig. 4D and 4E). As discussed below, this contrasts with the reduction in the rate of duplex unwinding and translation in response to eIF4A-R362Q, which potently inhibits translation at much lower concentrations than the concentration needed to appreciably reduce the overall rate of duplex unwinding (Fig. 3I and 3J). Together, our data show that inhibiting eIF4A or the recruitment of the eIF4F complex strongly reduces the rate of helicase activity in the cell-free extract. However, roughly 50% of the apparent helicase activity remains in the cell-free extract. This activity can remove the secondary structure located in the 5’ UTR of the mRNA reporter but is not sufficient for translation (Fig. 3 and Fig. 4).

### RNA duplex unwinding in nuclease-treated cell-free extracts is regulated by the m^7^G cap but not the poly (A) tail

Monitoring the rate of translation in the cell-free extracts reflects multiple rounds of translation. This makes it difficult to establish the relative role of the m^7^G cap and poly(A) tail in promoting the first round of initiation versus the subsequent 40S subunit binding events. Subsequent 40S binding events can include the binding of 40S subunits from the non-translating ribosome pool and/or the 40S subunits recycled from the mRNA following translation termination. We anticipate that our fluorescent helicase assay reflects the first eIF4F/40S subunit binding event on the mRNA reporter. This assumes that the fluorescent reporters will not reanneal to the mRNA following strand separation in the cell-free extract. While we are not able to directly measure this, we feel that this is reasonable given the fact that non-specific RNA binding proteins are present in the lysate and the addition of a competitor strand (used in our reconstitution assay) does not change the rate of duplex unwinding (data not shown). To determine the relative contributions of the m^7^G cap and poly(A) tail on the recruitment of the translation machinery in the first round of translation, we determined the rate of duplex unwinding on combinations of mRNAs in the absence (−) or presence (+) of a m^7^G cap (c), poly(A) tail (p), or both modifications (Fig. 5A).

**Figure 5.**
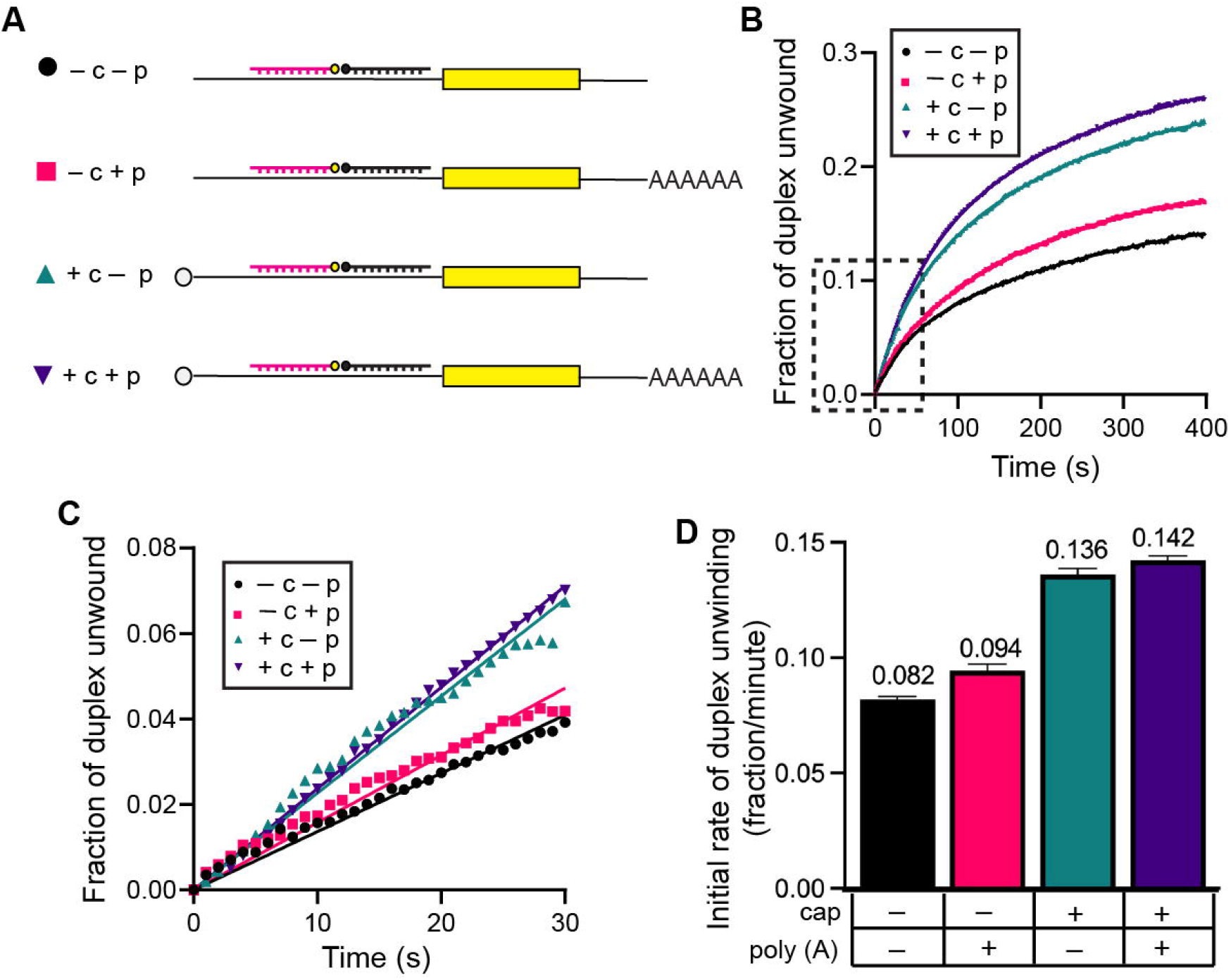
RNA duplex unwinding in nuclease-treated cell-free extracts is regulated by m^7^GTP cap but not poly(A) tail. **(A)** Schematic representation of the reporter constructs used in this set of experiments, with each combination of the m^7^GTP and poly(A) tail modifications to the mRNA. **(B)** The average trace of three independent time courses is shown for the fraction of duplex unwound in cell-free extracts in the presence of each mRNA reporter construct shown in (A). **(C)** A magnified view of the dashed region of the unwinding time course (B) is shown. Linear fits to the initial portion of the unwinding data for each mRNA reporter. **(D)** The initial rates of duplex unwinding (fraction unwound per min) determined by slope of the line in (C). Error bars for all data indicate the standard errors of the mean.

Even with very low background translation in the absence of the m^7^G cap and poly(A) tail (Fig. 2C), we observe an appreciable increase in the fraction of duplex unwound over time upon addition of the −c −p mRNA reporter to the cell-free extract (Fig. 5B). The initial rate of duplex unwinding per min for this mRNA (−c −p) is calculated to be 0.082 ± 0.001 (Fig. 5C, 5D, and Table 1). We note that the rate of unwinding is very similar to that observed for the +c +p mRNA reporter in the presence of eIF4A/4F inhibitors (Fig. 3, Fig. 4, and Table 1). The addition of the poly(A) tail (−c +p) does not appreciably increase the fraction of duplex unwound compared to the −c −p mRNA (Fig. 5B). Accordingly, the −c +p construct has a calculated initial rate of duplex unwinding of 0.094 ± 0.003 (Fig. 5C, 5D and Table 1). The inability of the poly(A) tail to promote duplex unwinding is consistent with its negligible effect on the rate of translation in the absence of m^7^G cap (Fig. 2C). Compared to the −c −p construct, we observe an appreciable increase in the fraction of duplex unwound over time in the presence of the m^7^G cap, +c −p (Fig. 5B). Calculating the initial rate of duplex unwinding shows that the initial fraction of duplex unwound per minute is increased by 1.6-fold from 0.082 ± 0.001 for −c −p to 0.136 ± 0.003 for +c −p (Fig. 5C, 5D and Table 1). In the presence of the m^7^G cap and poly(A) tail (+c +p), we do not observe any additional stimulation of duplex unwinding compared to the +c −p construct (Fig 5B). Our data therefore show that the m^7^G cap structure is the primary modification responsible for promoting duplex unwinding in our cell-free extract system. The poly(A) tail synergizes with the m^7^G cap for translation (Fig. 2), but we observe very little, if any, stimulation of duplex unwinding by the addition of the poly(A) tail (Fig. 5 and Table 1).

## DISCUSSION

In this study, we have developed a real-time fluorescent duplex unwinding assay to precisely monitor RNA helicase activity in the 5’ UTR of a mRNA using a translationally active cell-free extract. To this end, we engineered binding sites for two previously characterized fluorescently modified reporter strands into the 5’ UTR of a full-length mRNA that encodes NLuc. Introducing the mRNA into a cell-free extract generated from HeLa cells enables the parallel monitoring of duplex unwinding and translation of the mRNA using the same conditions (Fig. 1). Previous work has shown that the eIF4A helicase is required for duplex unwinding in various reconstituted systems (5,45,46). Accordingly, it has generally been assumed that the helicase activity of eIF4A is required for fundamental duplex unwinding on all mRNAs that contain secondary structure in their 5’ UTR. Nevertheless, additional helicase proteins including DDX3/ded1p and DHX29 have been shown to regulate the translation of specific mRNAs, especially those that are predicted to possess significant stable secondary structure in their 5’ UTRs (16–19,24). It is therefore unclear to what extent secondary structure in the 5’ UTR of mRNA is remodelled by eIF4A versus other helicase proteins. Recently, eIF4A has been shown to promote mRNA recruitment to the 40S subunit in a reconstituted purified system independently of its helicase activity (42). Moreover, eIF4A has been shown to promote translation on all mRNAs irrespective of their secondary structure (15). Thus, eIF4A may use its ATP binding activity to modulate the conformation of the 40S subunit for binding mRNA in addition to unwinding secondary structure.

To precisely determine the proportion of duplex unwinding activity attributable to eIF4A in HeLa cell-free extracts, we specifically inhibited eIF4A activity by adding Hippuristanol or a recombinantly expressed and purified dominant negative eIF4A mutant (eIF4A-R362Q). Consistent with previous reports, we find that both methods of inhibiting eIF4A strongly reduces mRNA translation of a luciferase reporter in HeLa cell-free extracts (Fig. 3E and 3J). Despite the addition of a concentration of Hippuristanol that completely inhibits luciferase translation, we were surprised to observe only a partial inhibition (~40%) of duplex unwinding (Fig. 3). To better understand the relationship between eIF4A-dependent duplex unwinding and translation, we titrated an increasing concentration of purified eIF4A-R362Q into the cell-free extract. Our data show that a concentration of eIF4A-R362Q that almost completely inhibits translation (1.5 μM) only reduces the rate of duplex unwinding of our reporter mRNA by ~10% (Fig. 3I and 3J). This suggests that robust duplex unwinding in the cell-free extract is not sufficient for promoting translation initiation in the presence of eIF4A-R362Q. In addition to its ability to unwind secondary structure, our data support a model where eIF4A also promotes translation by a helicase-independent mechanism. This helicase-independent mechanism may include the ATP-dependent, but helicase-independent, activity of eIF4A in promoting mRNA recruitment to the 40S subunit (42).

It is striking that even at the highest concentration of eIF4A-R362Q, a considerable rate of duplex unwinding on the mRNA reporter is still observed in the cell-free extract. Hippuristanol or eIF4A-R362Q may not prevent mRNA recruitment to form the 48S complex. The mammalian DDX3 and DHX29 helicases have been proposed to unwind mRNA secondary structure via their interactions with eIF4G and the 40S subunit (16,19,20). In the presence of Hippuristanol and eIF4A-R362Q, it is therefore possible that DDX3/ded1p and DHX29 may be responsible for some of the observed eIF4A-independent duplex unwinding activity. To prevent mRNA recruitment to eIF4F and the 48S complex, we titrated an increasing concentration of a competitor eIF4E protein (eIF4E-W73L) into the cell-free extract. Our data reveal a concentration dependent reduction in the rate of translation and the rate of duplex unwinding in response to eIF4E-W73L addition (Fig. 4). In contrast to eIF4A-R362Q, the observed reduction in the rate of duplex unwinding and the reduced rate of translation more closely matches each other as the eIF4E-W73L concentration is increased. This observation is consistent with eIF4E-W73L functioning as a competitive inhibitor to eIF4F. Importantly, the inhibition of duplex unwinding by eIF4E-W73L still appears to reach a plateau that reveals an appreciable amount of residual duplex unwinding activity that is likely eIF4F-independent (Fig. 4). We do note that this plateau appears to be slightly below that for eIF4A-R362Q (Fig. 3, Fig. 4 and Table 1). Whether the difference in these plateaus reflect the additional duplex unwinding activities of DDX3 and DHX29 remains an open question. Taken together, our data reveal a substantial amount of duplex unwinding in cell-free extracts is independent of the eIF4A helicase. The identity of the other helicase(s) responsible for promoting this duplex unwinding activity is not clear but may include DDX3 and DHX29.

Global monitoring of RNA structure has revealed that there are vastly fewer structured mRNA regions found in live yeast and mammalian cells than expected (25). The absence of extensive mRNA secondary structure in live cells was partly attributed to energy-dependent processes, such as RNA helicases (25). Importantly, the authors propose that translation is not likely to be the dominant process for unfolding RNA structure in live cells since the absence of secondary structure in mRNAs was not significantly different between coding regions and 5’ and 3’ UTRs (25). Our data are consistent with the presence of robust helicase activity being present in cells to remove secondary structure independently of the translation machinery. Nevertheless, eIF4A-dependent unwinding is observed in our cell-free extract, the activity of which is required for efficient translation. While the cell-free extract displays eIF4A-dependent and eIF4A-independent duplex unwinding activity, we recognize that the process of generating our cell-free extracts disrupts the cellular architecture. It is possible that this process may expose a reporter mRNA to helicases that it would not normally be physiologically exposed to in a live cell. While the physiological significance can be debated, cell-free extracts are extensively used to study the role of mRNA secondary structure in regulating translation and other aspects of mRNA lifecycle. Thus, our discovery of extensive eIF4A-independent duplex unwinding in cell-free extract will be relevant for interpreting future studies and for re-evaluating the interpretation of previous studies.

We have also used our duplex unwinding assay to determine the relative contributions of the m^7^G cap and poly(A) tail on the recruitment of the translation machinery in the first round of translation (Fig. 5A). Our data demonstrate that the m^7^G cap increases the rate of duplex unwinding by 1.6-fold, but no further increase in the rate of duplex unwinding was apparent from the presence of a poly(A) tail. Importantly, the poly(A) tail strongly promotes the rate of translation in our cell-free extract, indicating that our system possesses faithful regulation of translation by the poly(A) tail. Our data are therefore consistent with the poly(A) tail promoting a translation step after the initial recruitment of mRNA to the eIF4F and/or 48S complex. Nevertheless, we highlight that the effect of the poly(A) tail on translation in our system is measured over a 30-minute period, compared to the first few seconds/minutes for duplex unwinding. It is therefore possible that a very slow recycling step exists in our cell-free extract that releases PABP from any remaining mRNA fragments that are generated during nuclease digestion. To minimize the likelihood this possibility, we preincubated the cell-free extract for 10 minutes at 30 °C to ensure that the translation machinery is active and recycled from any nuclease digested mRNA fragments (Fig. S1B). We therefore feel that our data support a previously proposed model whereby the poly(A) tail promotes the recycling of ribosomes, which will promote the rate of ribosome reentry into translation initiation (47).

In this study, we have shown that our fluorescent duplex unwinding assay can be used to study duplex unwinding in a translationally competent cell-free extract system. We have used this method to reveal robust eIF4A-dependent and eIF4A-independent duplex unwinding occurs on a reporter mRNA. Unexpectedly, we show that roughly 50% of the duplex unwinding activity on a reporter mRNA is promoted by an eIF4A-independent mechanism. In future, it will be important to identify the helicase(s) that promote this eIF4A-independent duplex unwinding. We highlight the fact that our cell-free extract duplex unwinding assay can successfully measure the inhibitory nature of the eIF4A specific inhibitor Hippuristanol. Thus, we anticipate that other small molecule inhibitors could be tested for potential helicase inhibition using our cell-free duplex unwinding assay. Alternatively, one could generate specific helicase-depleted HeLa cell-free extracts (39) to study their possible contribution to the observed eIF4A-independent duplex unwinding activity.

## EXPERIMENTAL PROCEDURES

### Recombinant proteins

Mutant proteins for human eIF4A (eIF4A-R362Q) and eIF4E (eIF4E-W73L) were generated by site directed mutagenesis and confirmed by sequencing. Recombinant proteins were expressed and purified from *E. coli* BL21 (DE3), as described previously for wild type eIF4A and eIF4E (5,28).

### mRNA reporters

RNAs were transcribed by T7 RNA polymerase using a protocol described previously (27). The DNA template was generated by PCR of a reporter plasmid that contains a beta globin 5□ UTR, fluorescent beacon binding sequences, a NLuc ORF, and a short 3□ UTR (Fig. S3). The PCR primers include a T7 promoter in the forward primer and a reverse primer +/− a 50 nt poly(A) sequence (Fig. S3). The PCR product was verified to be free of aberrant bands by agarose gel electrophoresis. PCR templates were purified by phenol-chloroform extraction followed by ethanol precipitation. The RNA was transcribed with T7 polymerase and purified by phenolchloroform extraction followed by ethanol precipitation with ammonium acetate. Free nucleotides were removed from the RNA using Sephadex G25 resin (GE Healthcare) and RNA integrity and purity was verified by denaturing urea-polyacrylamide gel electrophoresis. The Nluc reporter mRNAs were capped using the Vaccinia Capping System (New England Biolabs) according to the manufacturer’s instructions (29).

Fluorescent reporter RNA oligonucleotides are chemically synthesized, modified, and HPLC purified by Integrated DNA Technologies (IDT). The reporter strand is modified with cyanine 3 (Cy3) on its 5’-end and the quenching strand is modified with a spectrally paired black hole quencher (BHQ) on its 3’-end (27).

### HeLa cell-free extract

HeLa S3 cells were grown in suspension as previously described with minor modifications (29). Two litres of exponentially growing cells were harvested at a density between 0.25×10^6^ and 0.30×10^6^ cells/ml. Cells were pelleted at 700 rcf for 10 min using a JLA-8.1000 rotor in a Beckman Coulter Avanti JXN-26 centrifuge at 4 °C. The cell pellets were pooled, washed three times with 10 ml ice-cold PBS (HyClone), and approximate volume of cell pellet recorded after the final wash. The cell pellet was resuspended in a 2-fold vol/vol of lysis buffer [10 mM Hepes (pH 7.5), 10 mM Potassium acetate, 0.5 mM Magnesium acetate, 5 mM DTT and one cOmplete, mini, EDTA-free protease inhibitor cocktail tablet (Roche)] and incubated for 5 min on ice. Cells were lysed for 3 × 20 sec at 4 °C using a Virtis homogenizer, transferred to Eppendorf tubes and centrifuged at 13,000 × g in a microfuge for 5 min at 4 °C. Lysates were then treated with 150 U/ml micrococcal nuclease (Fermentas) in the presence of 1 mM CaCl_2_ for 20 min at 18 °C before inactivation with 2 mM EGTA. Lysate was typically found to be 30–50 ODU/ml (A280 nm) and were flash frozen in liquid nitrogen and stored at −80°C.

### In vitro luciferase translation assay

Luciferase translation assays were carried out in 70 μl reactions containing 40% nuclease treated HeLa lysate and the following buffer components: 90 mM potassium acetate, 45 mM KCl, 2 mM magnesium acetate, 0.4 U/μl RNasin (Promega), 60 μM amino acid mixture (Promega), 50 μM Spermidine, 16 mM Hepes pH 7.5, 20 mM creatine phosphate, 40 μg creatine phosphokinase, 0.8 mM ATP-Mg, 0.1 mM GTP-Mg. For each reaction, 7 μl 500 nM mRNA (50 nM final concentration unless other stated) was added in reaction buffer (20 mM Tris-Acetate pH 7.5, 2 mM Mg-Acetate pH 7.5, 0.2 mM DTT, 100 mM KCl, 10 % glycerol). The addition of 7 μl purified protein or inhibitor was also added in reaction buffer. Prior to adding the mRNA reporter, reactions are incubated in the absence or presence of purified protein/inhibitors at 30 °C for 10 minutes. This preincubation time was found to be important in maintaining a linear rate of translation for the first 30 minutes (Fig. S1B) Following preincubation, a final concentration of 50 nM mRNA is added, and the translation reaction is incubated at 30 °C for 30 minutes (or as indicated). Each 70 μl reaction is separated into three individual 20 μl reactions and 100 μl of Renilla Luciferase assay substrate (Promega) is added to each reaction. Renilla Luciferase assay substrate was found to provide appropriate signal for the Nano Luciferase reporter. After mixing, 100 μl of each reaction was transferred to a 96 well OptiPlate (Perkin Elmer) and measured using a Victor X5 Multilabel Plate Reader (Perkin-Elmer). Translation data were normalized to a control reaction containing the same buffer without mRNA reporter. Translation is quantified as the average of at least three independent experiments. Error bars represent the standard error of the mean.

### Duplex unwinding assay

A reporter RNA (5□-Cy3-GUUUUUUAAUUUUUUAAUUUUUUC-3□) and quencher RNA (5□GGCCCCACCGGCCCCUCCG-BHQ-3□) were annealed to the mRNA template RNA (1:1:1 ratio) at 500 nM by heating to 80°C and slow cooling to room temperature in unwinding buffer [20 mM Tris-acetate (pH 7.5), 2 mM magnesium acetate, 100 mM potassium acetate, and 0.2 mM DTT]. The final annealed reporter mRNA is diluted to 250 nM on ice using unwinding buffer. Duplex unwinding reactions are performed at 30 °C in a 50 μl cuvette (Starna) using a Fluorolog-3 spectrofluorometer, as describe previously with modifications (27). To calibrate the maximum fluorescence for each experiment, a mock unwinding assay is prepared in the absence of the BHQ-modified oligonucleotide. In addition to determining the maximum fluorescence, this mock assay enables one to determine if there is appreciable photobleaching in the lysate. For unwinding reactions, 14 μl annealed mRNA substrate (250 nM), 21 μl reaction buffer and 7 μl (20 mM ATP-Mg) is added to the cuvette and the baseline is monitored to verify the annealed substrate is stable. To initiate the reaction, 28 μl preincubated lysate is added to the cuvette (70 μl reaction total) and the reaction is monitored for 10 minutes.

The data were analyzed as described previously with minor modifications (27). Noise from the lamp intensity (R1) was removed from the data points (S1) using the formula S1/R1. The fraction of duplex unwound was calculated using the maximum fluorescence and the baseline fluorescence immediately measured after the addition of cell lysate. The baseline fluorescence is subtracted from the values for each time point and from the maximum fluorescence. The fraction of duplex unwound is calculated by dividing the baseline-corrected fluorescence values at each time point by the corrected maximum fluorescence. The initial rate of duplex unwinding was determined by a linear fit of the initial portion of the unwinding reaction and converted to the fraction of duplex unwound per minute. Each unwinding reaction is quantified as the average of at least three independent experiments. Error bars represent the standard error of the mean.

## Data availability

Data is presented within the manuscript and plasmids and other reagents are available for academic purposes upon request.

## Supporting information

This article contains supporting information.

## Acknowledgements

We wish to thank the Fraser laboratory for many insightful comments and critical reading of the manuscript.

## Author Contributions

M.H.O. and C.S.F. conceived and designed the experiments and led the project. M.H.O carried out all experiments and both authors wrote and edited the manuscript.

## Funding and additional information

This work was supported by NIH grant R01 GM092927 to CSF and an NIH T32 Training Program in Molecular and Cellular Biology (T32GM007377) to M.H.O. The content is solely the responsibility of the authors and does not necessarily represent the official views of the National Institutes of Health.

## Conflict of interest

The authors declare that they have no conflicts of interest with the contents of this article.

## Notes

### Competing Interest Statement

The authors have declared no competing interest.

