## Supporting Information for "Monitoring RNA restructuring in a human cell-free extract reveals eIF4A-dependent and eIF4A-independent unwinding activity"

### Supplementary Figure 1

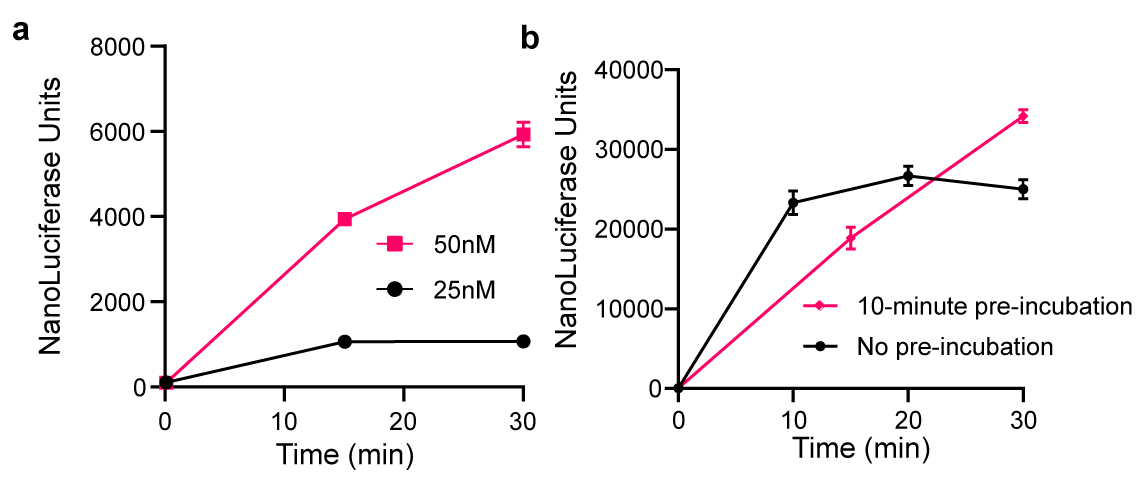

#### Figure S1. Optimization of translation conditions to ensure linearity of protein synthesis in nuclease treated cell-free extract. (a) Line graph depicting luciferase translation of mRNA reporter measured over 30 min at an incubation temperature at 30 °C. *In vitro* translation reactions in lysate programmed with 25 nM or 50 nM mRNA reporter. (b) Line graph demonstrating the importance of a 10‑minute pre-incubation step prior to initiating protein synthesis. *In vitro* translation reactions in lysate programmed with 50 nM reporter over a time course of 30 minutes. All data are presented as means of three independent experiments ± SEM.

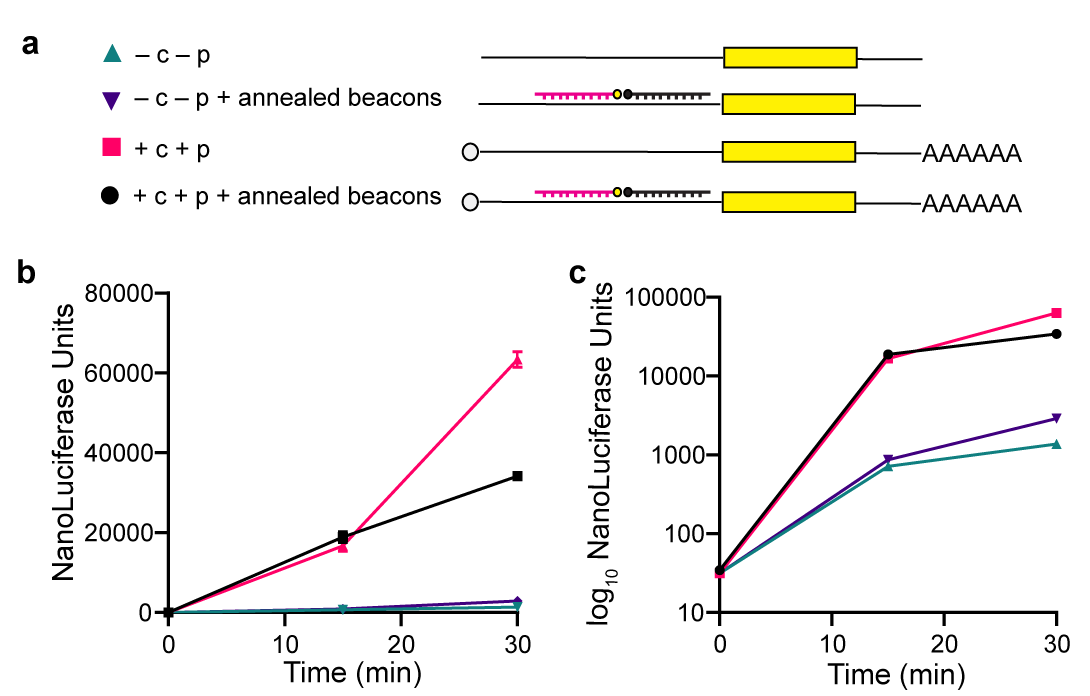

### Supplementary Figure 2

### Figure S2. Effect of annealed beacons on protein synthesis. (a) Schematic representation of the reporter constructs used in this set of experiments indicating the combination of m^7^G cap and poly(A) tail and fluorescent beacons annealed to the mRNA 5′ UTR. (b) Line graph depicting luciferase translation of mRNA reporter measured over 30 min at an incubation temperature at 30 °C. *In vitro* translation reactions in lysate programmed with 50 nM mRNA reporter in the absence or presence of m^7^G cap, poly(A) tail, and annealed fluorescent beacons. (c) Line graph using data from (b) but with the Y-axis transformed to a log_10_ scale. All data are presented as means of three independent experiments ± SEM.

**Supplementary Figure 3**

Dual-assay mRNA Reporter Sequence

Parental Plasmid (pUC57) that includes the Globin 5'UTR; 24nt-CY3; 19nt-BHQ, NLuc ORF.

GGTACC: KpnI restriction site.

T7 promoter between KpnI and BamHI sites.

GGATCC: BamHI restriction site.

B-globin 5' UTR.

CCATGG: NcoI restriction site.

Helicase Double Reporter: 24nt Cy3 binding site, 19nt BHQ binding site

CTGCAG: PstI cut site

Nanoluciferase reporter

AAGCTT: HindIII restriction site

Reporter template sequence:

GGTACCTAATACGACTCACTATA**G**GGGGATCCacatttgcttctgacacaactgtgttcactagcaacctcaaacagacaccCTGCAGGAAAAAATTAAAAAATTAAAAAACTCGGAGGGGCCGGTGGGGCCACCATGGTCTTCACACTCGAAGATTTCGTTGGGGACTGGCGACAGACAGCCGGCTACAACCTGGACCAAGTCCTTGAACAGGGAGGTGTGTCCAGTTTGTTTCAGAATCTCGGGGTGTCCGTAACTCCGATCCAAAGGATTGTCCTGAGCGGTGAAAATGGGCTGAAGATCGACATCCATGTCATCATCCCGTATGAAGGTCTGAGCGGCGACCAAATGGGCCAGATCGAAAAAATTTTTAAGGTGGTGTACCCTGTGGATGATCATCACTTTAAGGTGATCCTGCACTATGGCACACTGGTAATCGACGGGGTTACGCCGAACATGATCGACTATTTCGGACGGCCGTATGAAGGCATCGCCGTGTTCGACGGCAAAAAGATCACTGTAACAGGGACCCTGTGGAACGGCAACAAAATTATCGACGAGCGCCTGATCAACCCCGACGGCTCCCTGCTGTTCCGAGTAACCATCAACGGAGTGACCGGCTGGCGGCTGTGCGAACGCATTCTGGCGTAAAAGCTT

Forward primer used to PCR template:

5′-CGTTGTAAAACGACGGCCAG-3′

Reverse primer used to PCR template:

5′-(T)_50_ACCCCAGGCTTTACACTTTATGC-3′

Reverse primer is complementary to the pUC57 sequence downstream of the reporter sequence and adds a 100 nt 3′ UTR sequence from the pUC57 plasmid.
