## Supplementary material for "Monitoring RNA restructuring in a human cell-free extract reveals eIF4A-dependent and eIF4A-independent unwinding activity": Table 1

TABLE 1. Summary of duplex unwinding and translation data.

| **Condition** | **Initial rate of duplex Unwinding**  **(Fraction/min)** | **NLuc translation**  **(Units)** |
| --- | --- | --- |
| 2 mM ATP | 0.189 ± 0.001 | 48202 ± 3022 |
| 10 mM AMP-PNP | 0.024 ± 0.001 | 170 ± 55 |
| 0.3 % DMSO | 0.162 ± 0.002 | 11230 ± 1105 |
| 3 μM Hippuristanol  (in 0.3 % DMSO) | 0.104 ± 0.001 | 94 ± 2 |
| 0 μM eIF4A-R362Q | 0.159 ± 0.003 | 7814 ± 532 |
| 1.5 μM eIF4A-R362Q | 0.144 ± 0.003 | 649 ± 42 |
| 3.0 μM eIF4A-R362Q | 0.117 ± 0.001 | 383 ± 27 |
| 6.0 μM eIF4A-R362Q | 0.102 ± 0.001 | 333 ± 23 |
| 0 μM eIF4E-W73L | 0.167 ± 0.003 | 9449 ± 953 |
| 0.5 μM eIF4E-W73L | 0.133 ± 0.002 | 8786 ± 217 |
| 1.0 μM eIF4E-W73L | 0.095 ± 0.001 | 4626 ± 185 |
| 2.0 μM eIF4E-W73L | 0.080 ± 0.001 | 3030 ± 101 |
| m^7^G cap – poly(A) – | 0.082 ± 0.001 | 1049 ± 53 |
| m^7^G cap – poly(A) + | 0.094 ± 0.003 | 2338 ± 56 |
| m^7^G cap + poly(A) – | 0.136 ± 0.003 | 15205 ± 452 |
| m^7^G cap + poly(A) + | 0.142 ± 0.002 | 29930 ± 1888 |
